# Improving prediction of drug-target interactions based on fusing multiple features with data balancing and feature selection techniques

**DOI:** 10.1101/2022.12.07.519302

**Authors:** Hakimeh Khojasteh, Jamshid Pirgazi

## Abstract

Predicting drug-target interaction (DTI) is an important research area in the field of drug discovery. It means identifying the interaction between chemical compounds and protein targets. Wet lab experiments to explore these interactions are expensive as well as time-consuming. On the contrary, a dry lab environment focusing more on computational methods of interaction prediction can be helpful to limit the search space for the wet lab experiments and give clues before developing a new medicine. This paper proposes a novel drug-target interaction prediction method called SRX-DTI. First, we extract various descriptors from protein sequences, and the drug is encoded as FP2 fingerprint. Besides, we present the One-SVM-US technique to deal with imbalanced data. We also developed the FFS-RF algorithm, a forward feature selection algorithm, and coupled it with a random forest (RF) classifier to maximize the predictive performance. This feature selection algorithm removes the irrelevant features to obtain the best optimal features. Finally, the balanced dataset with optimal features is given to the XGBoost classifier to identify DTIs. The experimental results demonstrate that our proposed approach SRX-DTI achieves significantly higher performance than other existing methods in predicting DTIs. The experimental results demonstrate that our proposed approach SRX-DTI achieves significantly higher performance than other existing methods in predicting DTIs. The datasets and source code are available at: https://github.com/Khojasteh-hb/SRX-DTI.

## 1. Introduction

The main phase in the drug discovery process is to identify interactions between drugs and targets (or proteins), which can be performed by in vitro experiments. Identifying drug-target interaction plays a vital role in drug development that aims to identify new drug compounds for known targets and find new targets for current drugs [1, 2]. The expansion of the human genome project has provided a better diagnosis of disease, early detection of certain diseases, and identifying drug-target interactions (DTIs) [3]. Although significant efforts have been done in the previous years, only a limited number of drug candidates have been permitted to reach the market by the Food and Drug Administration (FDA) whereas the maximum number of drug candidates have been rejected during clinical verifications, due to side effects or low efficacy [4]. Moreover, the cost of a new chemistry-based drug is often 2.6 billion dollars, and it takes typically 15 years to finish the drug development and approval procedure. This issue has been changing into a bottleneck to identify the targets of any candidate drug molecules [2, 5]. The experiment-based methods involve high cost, time-consuming, and small-scales limitations that motivate researchers to constantly develop computational methods for the exploitation of new drugs [2, 6, 7]. On the other side, availability of online databases in this area, such as KEGG [8, 9], DrugBank [10], PubChem[11], TTD [12, 13], and STITCH [14] have been influencing Machine Learning (ML) researchers to develop high throughput computational methods.

Besides developing computational methods in DTI prediction, studying protein-protein interactions (PPIs) has become a top priority for drug discovery, especially due to the SARS-CoV-2 pandemic[15–17]. Proteins are responsible for various essential processes in vivo via interactions with other molecules. Proteins are the building blocks of all diseases, and their malfunctioning is often responsible for diseases, making them crucial targets for the drug discovery process[18, 19]. The studies suggest that abnormal activity can support the development of life-threatening diseases like cancer. As a result, developing computational methods for identifying the critical proteins in PPIs has become an important branch of drug discovery and treatment development[19, 20].

The prior methods in DTI prediction can be mainly categorized into similarity-based methods and feature-based methods. In similarity-based methods, similar drugs or proteins are considered to find similar interaction patterns. These methods use many different similarity measures based on drug chemical similarity and target sequence similarity to identify drug-target interaction [21–23]. Feature-based methods consider drug–target interaction prediction as a binary classification problem and different classification algorithms such as Support Vector Machine (SVM) [24], random forest [25], rotation forest [26, 27], XGBoost [28], and deep learning [29, 30] have been employed to identify new interactions.

For drug-target prediction, Mousavian et al. applied a support vector machine to predict novel targets on features extracted from the Position Specific Scoring Matrix (PSSM) of proteins along with the molecular substructure fingerprint of drugs [24]. Another ML method based on the random forest is LRF-DTIs presented by Shi et al. [25]. In this method, the pseudo-position specific scoring matrix (PsePSSM) and the FP2 molecular fingerprint were used to extract the features from proteins and drugs and Lasso dimensionality reduction was used to reduce the dimension of the extracted features. The authors employed Synthetic Minority Oversampling Technique (SMOTE) method to deal with unbalanced data. Then, they predicted DTIs by the random forest (RF) classifier using feature vectors. Wang et al. proposed an approach namely RFDT: A Rotation Forest-based Predictor that used a feature vector as PSSM descriptor for protein and the drug molecules fingerprint [27]. Recently, another method based on Rotation Forest has been developed by Wang et al. [26] is RoFDT that combines feature-weighted Rotation Forest (FwRF) with a protein sequence. They also encode protein sequences as PSSM and extracted their features utilizing PsePSSM. Finally, the combination of those features with drug structure fingerprints was fed into the FwRF classifier to predict DTIs.

Moreover, Mahmud et al. [28] proposed a computational model, called iDTi-CSsmoteB for the identification of DTIs. They utilized PSSM, amphiphilic pseudo amino acid composition (AM-PseAAC), and dipeptide PseAAC descriptors to present protein. For drug chemical structure, they used molecular substructure fingerprint (MSF) which describes the existence of the functional fragments. Then, the oversampling SMOTE technique was applied to handle the imbalance of datasets, and the XGBoost algorithm as a classifier to predict DTIs.

With significant growth in the volume and variety of data, various platforms and libraries using deep learning have been developed such as DeepPurpose [29]and DeepDrug [30]. DeepPurpose [29] takes the SMILES format of the drug and amino acid sequence of protein as input and by a specific function, transformed it into a specific format. Finally, it is converted to a vector representation to use in the next steps. This library provides eight encoders using different modalities of compounds. DeepPurpose also provides utility functions to load a pre-trained model and predict new drugs and new targets. Another deep learning framework namely DeepDrug was proposed by Yin et al. [30]. Also, deep learning frameworks based on variants of graph neural networks such as graph convolutional networks (GCNs) [31], graph attention networks (GATs) [32, 33], gated graph neural networks (GGNNs) [34, 35] have developed for DTI prediction.

Here, to further improve prediction performance, we present SRX-DTI, a novel ML-based method for drug-target interaction prediction. First, we generate several descriptors of protein sequences including Amino acid composition (AAC), Dipeptide composition (DPC), Grouped amino acid composition (GAAC), Dipeptide deviation from expected mean (DDE), Pseudo amino acid composition (PseAAC), Pseudo-position-specific scoring matrix (PsePSSM), Composition of k-spaced amino acid group pairs (CKSAAGP), Grouped dipeptide composition (GDPC), and Grouped tripeptide composition (GTPC); and the drug is encoded as FP2 molecular fingerprint. Secondly, we present the One-SVM-US technique to make balancing, and the positive and negative samples are constructed by using the drug-target interactions information on the extracted features. Then, we perform the FFS-RF algorithm to select the optimal subset of features. Finally, after comparing different ML classifiers, the XGBoost classifier is chosen for predicting DTIs under the 5-Fold cross-validation. The performance evaluation is conducted by using AUROC, AUPR, ACC, SEN, SPE, and F1-score metrics. Area Under Receiver Operating Characteristic curve (AUROC) values of SRX-DTI on enzyme (EN), GPCR, ion channel (IC), and NR are 0.9920, 0.9880, 0.9788, and 0.9329 respectively. The results indicate that our proposed method significantly improved the prediction performance of DTIs in comparison with other existing methods.

The rest of the paper is organized as follows: Materials and methods section describes the detail of the gold standard datasets, feature extraction, data balancing, and feature selection, we utilized in this paper. In the Results and discussion section, performance evaluation and experimental results are provided. Finally, the Conclusions section summarizes the conclusions.

## 2. Materials and methods

In this study, we propose a novel method of drug-target interaction prediction, which is called SRX-DTI. At first step, drug chemical structures (SMILE format) and protein sequences (FASTA format) are collected from DrugBank and KEGG databases using their specific access IDs. In the next step, different feature extraction methods are applied to drug compounds and protein sequences to create a variety of features. Drug-target pair vectors are made based on known interactions and extracted features. Afterward, a balancing technique is utilized on DTI vectors to deal with imbalanced datasets, and drug–target features are selected through the FFS-RF to boost prediction performance. Finally, the XGBoost classifier is used on the balanced datasets with optimal features to predict DTIs. A schematic diagram of our proposed SRX-DTI model is shown in Fig. 1.

**Fig. 1.**
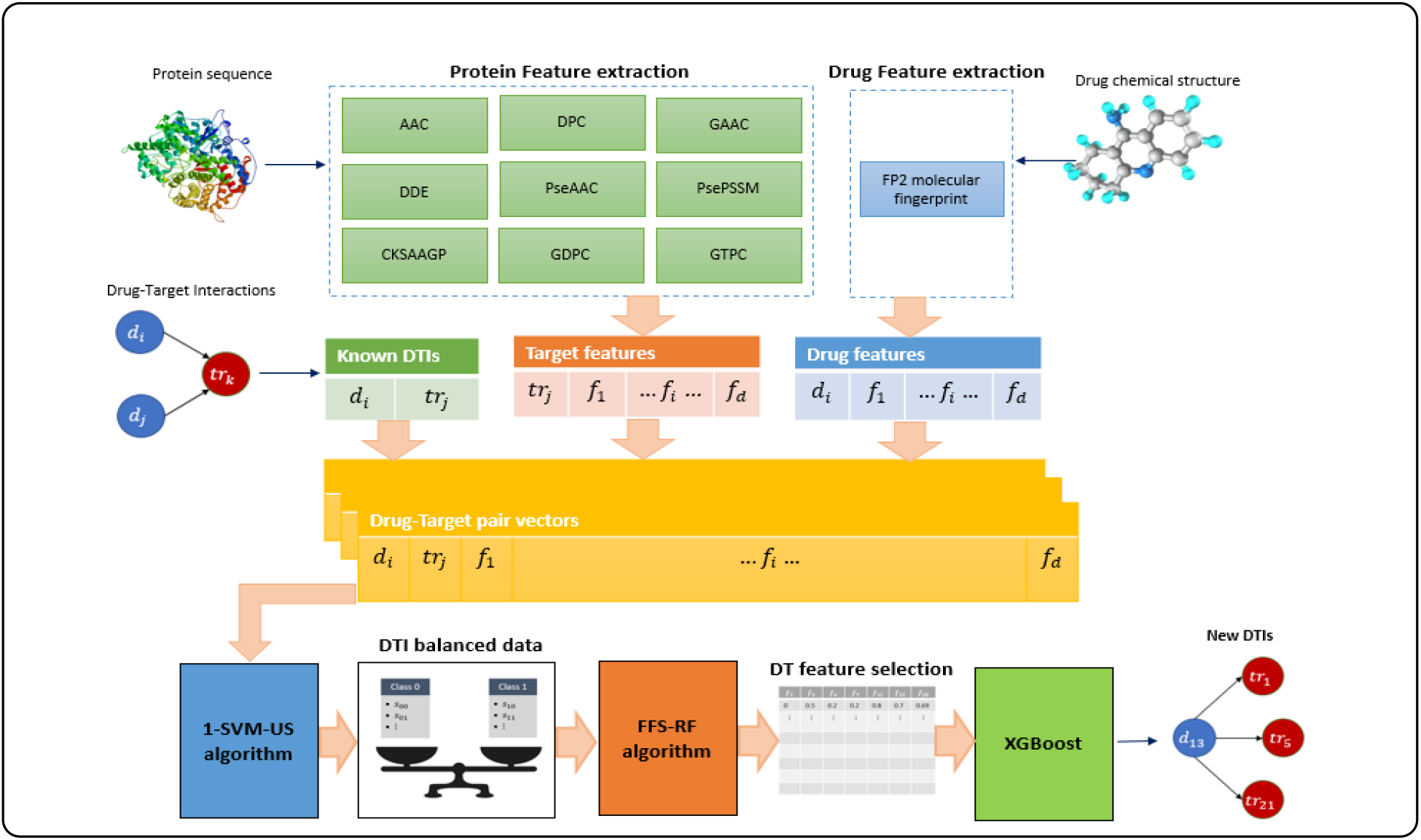
The workflow of the proposed model to predict drug-target interactions.

### 2.1 Drug–target datasets

In this research, four golden standard datasets, including enzymes (EN), G-protein-coupled receptors (GPCR), ion channel (IC), and nuclear receptors (NR) released by Yamanishi et al. [36] are explored as benchmark datasets to evaluate the performance of the proposed SRX-DTI method in DTI prediction. All these datasets are freely available from http://web.kuicr.kyoto-u.ac.jp/supp/yoshi/drugtarget/. Yamanishi et al. [36] extracted information about drug-target interactions from DrugBank [37], KEGG [8, 9], BRENDA [38] and SuperTarget [39]. The numbers of known interactions including enzymes, ion channels, GPCRs, and nuclear receptors are 2926, 1476, 635, and 90, respectively. A brief summary of these datasets is given in Table 1.

**Table 1.** Description of the gold standard datasets[36].

| Datasets | Drugs | Targets | Interactions |
| --- | --- | --- | --- |
| EN | 445 | 664 | 2926 |
| GPCR | 223 | 95 | 635 |
| IC | 210 | 204 | 1476 |
| NR | 54 | 26 | 90 |

## 3. Feature extraction methods

In order to better identify drug-protein interactions, it seems advantageous to extract different features from drugs and targets. This allows us to have more complete information about the known interactions and increase the detection rate. A brief summary of the ten groups of features is given in Table 2. Notice that there are two types of features. Drug related features and target related features in nine groups A, B, C, D, E, F, G, H, and I. In the following, these features are described, respectively. Based on drug and target descriptors, we constructed four subsets of features (AB, CD, EF, and GHI), which are given in Table 3. Also, notice that the drug features are coupled with singular target groups and these subsets.

**Table 2.** List of descriptors used in this study.

| Descriptor | Number of Features | Feature Type | Feature Group |
| --- | --- | --- | --- |
| Molecular fingerprint | 256 | drug |  |
| Amino acid composition (AAC) | 20 | target | A |
| Dipeptide composition (DPC) | 400 | target | B |
| Grouped amino acid composition (GAAC) | 5 | target | C |
| Dipeptide deviation from expected mean (DDE) | 400 | target | D |
| Pseudo amino acid composition (PseAAC) | 28 | target | E |
| Pseudo-position-specific scoring matrix (PsePSSM) | 220 | target | F |
| Composition of k-spaced amino acid group pairs (CKSAAGP) | 150 | target | G |
| Grouped dipeptide composition (GDPC) | 25 | target | H |
| Grouped tripeptide composition (GTPC) | 125 | target | I |

**Table 3.** Four subsets of features based on drug and target descriptors.

| Feature Combination | Number of Features |
| --- | --- |
| **AB**<br>(with drug features) | 676 |
| **CD**<br>(with drug features) | 661 |
| **EF**<br>(with drug features) | 504 |
| **GHI**<br>(with drug features) | 556 |

### 3.1 Drug features

For drug compounds, different types of descriptors can be defined based on various types of drug properties such as FP2, FP3, FP4 and MACCS [40–42]. Some studies showed that these descriptors are molecular structure fingerprints that effectively represent the drug [25, 43, 44]. In this study, the FP2 format fingerprint is used to present drug compounds. This molecular fingerprint of the drug was extracted through these steps:

Step 1: For each drug, molecular structure as mol format is downloaded from the KEGG database (https://www.kegg.jp/kegg/drug/) by using its drug ID.
Step 2: The OpenBabel Software (available from http://openbabel.org/) is downloaded and installed.
Step 3: The drug molecules with mol file format are converted into the FP2 format molecular fingerprint using the OpenBabel software. The FP2 format molecular fingerprint is a hexadecimal digit sequence of length 256 that is converted to a drug molecule 256-dimensional vector as a decimal digit sequence between 0 and 15.

### 3.2 Target features

#### A) Amino acid composition (AAC)

The amino acid composition [45] is a vector of 20 dimensions, which calculates the frequencies of all 20 natural amino acids (i.e. “*ACDEFGHIKLMNPQRSTVWY*”) as:

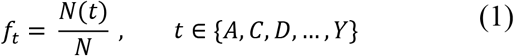

where *N*(*t*) is the number of amino acid type *t*, while *N* is the length of a protein sequence.

#### B) Dipeptide composition (DPC)

The Dipeptide Composition [46] gives 400 descriptors for protein sequence. It is calculated as:

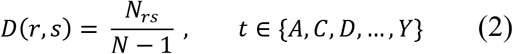

where *N_rs_* is the number of dipeptides represented by amino acid types *r* and *s* and *N* denotes the length of protein.

#### C) Grouped Amino Acid Composition (GAAC)

In the GAAC encoding [47], the 20 amino acid types are considered five classes according to their physicochemical properties. GAAC descriptor is the frequency of each amino acid group, which is calculated as:

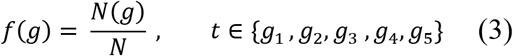

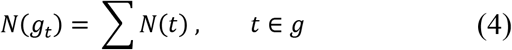

where *N*(*g*) is the number of amino acids in group *g*, *N*(*t*) is the number of amino acid type *t*, and *N* is the length of protein sequence.

#### D) Dipeptide Deviation from Expected mean (DDE)

The Dipeptide Deviation from Expected mean [46] is a feature vector, which is constructed by computing three parameters, i.e. dipeptide composition (*D_c_*), theoretical mean (*T_m_*), and theoretical variance (*T_v_*). These three parameters and the DDE are defined as follows. *D_c_*(*r*, *s*), the dipeptide composition measure for the dipeptide ‘*rs*’, is given as:

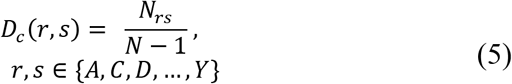

where *N_rs_* is the number of dipeptides represented by amino acid types *r* and *s* and *N* is the length of protein. *T_m_*(*r*, *s*), the theoretical mean, is given by:

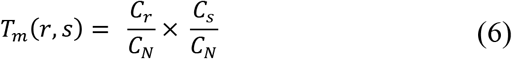

where *C_r_* is the number of codons, coding for the first amino acid, and *C_s_* is the number of codons, coding for the second amino acid in the given dipeptide ‘*r_s_*’ and *C_N_* is the total number of possible codons. *T_v_*(*r*, *s*), the theoretical variance of the dipeptide ‘*rs*’, is given by:

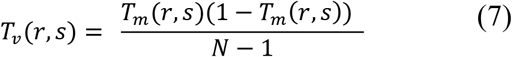

Finally, *DDE*(*r*, *s*) is calculated as:

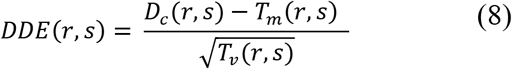

#### E) Pseudo Amino Acid composition (PseAAC)

To avoid completely losing the sequence-order information, the concept of PseAAC (pseudo amino acid composition) was proposed by Chou [48]; The idea of PseAAC has been widely used in bioinformatics including proteomics [49], system biology [50], such as predicting protein structural class [51], predicting protein subcellular localization [52], predicting DNA-binding proteins [53] and many other applications. In contrast with AAC which includes 20 components with each reflecting the occurrence frequency for One of the 20 native amino acids in a protein, the PseAAC contains a set of greater than 20 discrete factors, where the first 20 represent the components of its conventional amino acid composition while the additional factors are a series of rank-different correlation factors along a protein chain. According to the concept of PseAAC [48], any protein sequence formulated as a PseAAC vector given by:

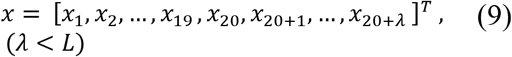

where *L* is the length of protein sequence, and λ is the sequence-related factor that choosing a different integer for, will lead to a dimension-different PseAAC. Each of the components can be defined as follows:

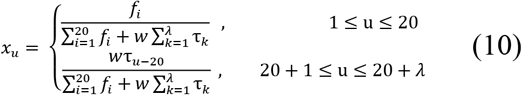

where *w* is the weight factor, and *f_i_* indicates the frequency at *i* – *th* AA in protein sequence. The τ_*k*_, the *k*-th tier correlation factor reflects the sequence order correlation between all the *k*-th most contiguous residues as formulated by:

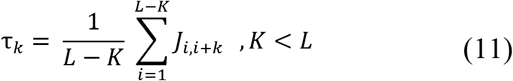

with

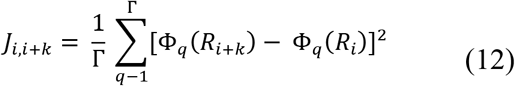

where Φ_*q*_(*R_i_*) is the *q*-th function of amino acid *R_i_*, and Γ is the total number of the functions considered. In this research, the protein functions which are considered, includes hydrophobicity value, hydrophilicity value, and side chain mass of amino acid. Therefore, the total number of functions Γ is 3.

In this study, λ is set to 1 and *W* is set to 0.05. The output characteristic dimensions of each target protein are 28 for the PseAAC descriptor.

#### F) Pseudo position specific scoring matrix (PsePSSM)

To represent characteristics of the amino acid (AA) sequence for protein sequences, the pseudo-position specific scoring matrix (PsePSSM) features introduced by Shen et al. [54] are used. The pseudo-position specific scoring matrix (PsePSSM) features encode the protein sequence’s evolution and information which have been broadly used in bioinformatics research [54–56].

For each target sequence P with L amino acid residues, PSSM is used as its descriptor proposed by Jones et al. [57]. The position-specific scoring matrix (PSSM) with a dimension of L×20 can be defined as:

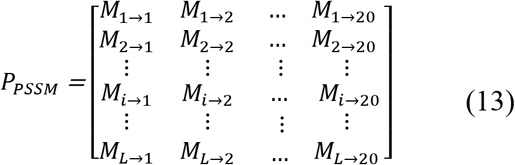

where *M_i,j_* indicates the score of the amino acid residue in the *i*th position of the protein sequence being mutated to amino acid type *j* during the evolution process. Here, for simplifying the formulation, it is used the numerical codes 1, 2,…, 20 to represent the 20 native amino acid types according to the alphabetical order of their single character codes. It can be searched using the PSI-BLAST [58] in the Swiss-Prot database. A positive score shows that the corresponding residue is mutated more frequently than expected, and a negative score is just the contrary.

In this work, the parameters of PSI-BLAST are set as the threshold of E-value equals 0.001, the maximum number of iterations for multiple searches equals 3, and the rest of the parameters by default. Each element in the original PSSM matrix was normalized to the interval (0, 1) using Eq. (14):

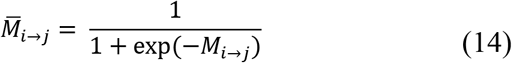

However, due to different lengths in target sequences, making the PSSM descriptor as a uniform representation can be helpful, one possible representation of the protein sample P is:

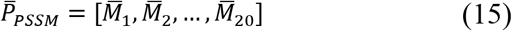

where T is the transpose operator, and

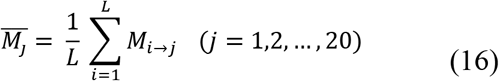

where *M*_*i*→*j*_ is the average score of the amino acid residues in the protein P changed to the *j*th amino acid residue after normalization, *M_j_* represents the average score of the amino acid residue in protein P being mutated amino acid type j during the process of evolution. However, if *P_PSSM_* of Eq. (13) represents the protein P, all the sequence-order information would be lost. To avoid complete loss of the sequence-order information, the concept of the pseudo amino acid composition introduced by Chou [59], i.e. instead of Eq. (11), we use position-specific scoring matrix (PsePSSM) to represent the protein P:

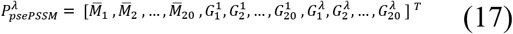

where

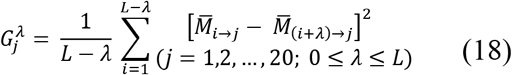

where 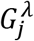 represents the correlation factor of the j - th amino acid and λ is the continuous distance along the protein sequence. This means that 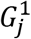 is the relevant factor coupled along the most continuous PSSM score on the protein chain of amino acid type *j*, 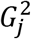 is the second closest PSSM score by coupling, and so on. Therefore, a protein sequence can be defined as Eq. (15) using PsePSSM and produces a 20 + 20 × *λ-dimensional* feature vector. In this study, *λ* is set to 10. The output characteristic dimension of each target protein is 220 for the PsePSSM descriptor.

#### G) Composition of k-spaced amino acid group pairs (CKSAAGP)

The Composition of k-Spaced Amino Acid Group Pairs (CKSAAGP) [60] defines the frequency of amino acid group pairs separated by any k residues (the default maximum value of k is set as 5). If *k* = 0, the 0-spaced group pairs are represented as:

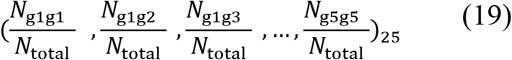

where the value of each descriptor indicates the composition of the corresponding residue group pair in a protein sequence. For a protein of length *P* and *k = 0, 1, 2, 3, 4* and *5*, the values of *N*_total_ are *P – 1, P– 2, P – 3, P – 4, P – 5* and *P – 6* respectively.

#### H) Grouped dipeptide composition (GDPC)

The Grouped Di-Peptide Composition encoding [60] is a vector of 25 dimensions, which is another variation of the DPC descriptor. It is defined as:

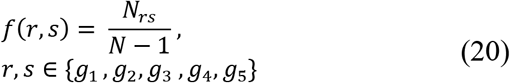

where *N_rs_* is the number of dipeptides represented by amino acid types *r* and *s* and *N* denotes the length of a protein.

#### I) Grouped tripeptide composition (GTPC)

The Grouped Tri-Peptide Composition encoding [60] is also a variation of the TPC descriptor, which generates a vector of 125 dimensions, defined as:

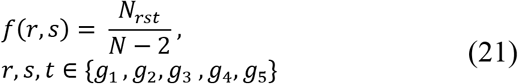

where *N_rst_* is the number of tripeptides represented by amino acid types *r*, *s* and *t. N* denotes the length of a protein.

## 4. Data balancing technique

The experiment datasets that we used in this study were highly imbalanced. Most of machine learning algorithms perform poorly on imbalanced datasets, and model will be completely biased toward majority instances, and it will ignore minority ones. Different techniques have been utilized to balance the imbalanced dataset, such as random undersampling [24, 61, 62], cluster undersampling [63, 64], and SMOTE technique [25, 28]. In this study, we developed a new undersampling algorithm namely One-SVM-US using One-class Support Vector Machine (SVM) to deal with imbalanced data. Also, the known DTIs are considered positive samples. For enzymes, ion channels, GPCRs, and nuclear receptors, the number of positives are 2926, 1476, 635, and 90, respectively. Our novel algorithm considers all of the possible interactions in four datasets as negative samples except the ones that have been known as positive. By performing the algorithm, it would result in a balanced dataset with equal numbers of positive and negative samples.

A One-Class Support Vector Machine (One-class SVM) [65], is a semi-supervised global anomaly detector. This algorithm needs a training set that contains only one class. The One-SVM-US technique based on One-class SVM considers all possible combinations of drug and target by discarding those that are positive samples. This algorithm uses a hypersphere to encompass all of the instances instead of using a hyperplane to separate two classes of samples. We apply the RBF kernel for SVM. The setting for the parameter γ was investigated, which was the simple heuristic γ=1/no. of data points. To compute the outlier score, first, compute the maximum value of the decision function:

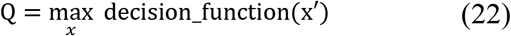

where *x* refers to the vector of scores. Then, we obtained the outlier score as follows:

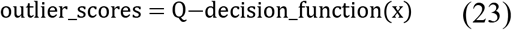

Then, the outlier scores are sorted in ascending and the *n_minority_* samples are selected from the sorted list. The final data is constructed from the combination of the minority class from the original experimental dataset and the majority class chosen by the proposed method. Even though, we would like to mention that Algorithm 1 performs effectively to make balanced datasets.

### Algorithm 1: Undersampling by One-class SVM-(One-SVM-US)

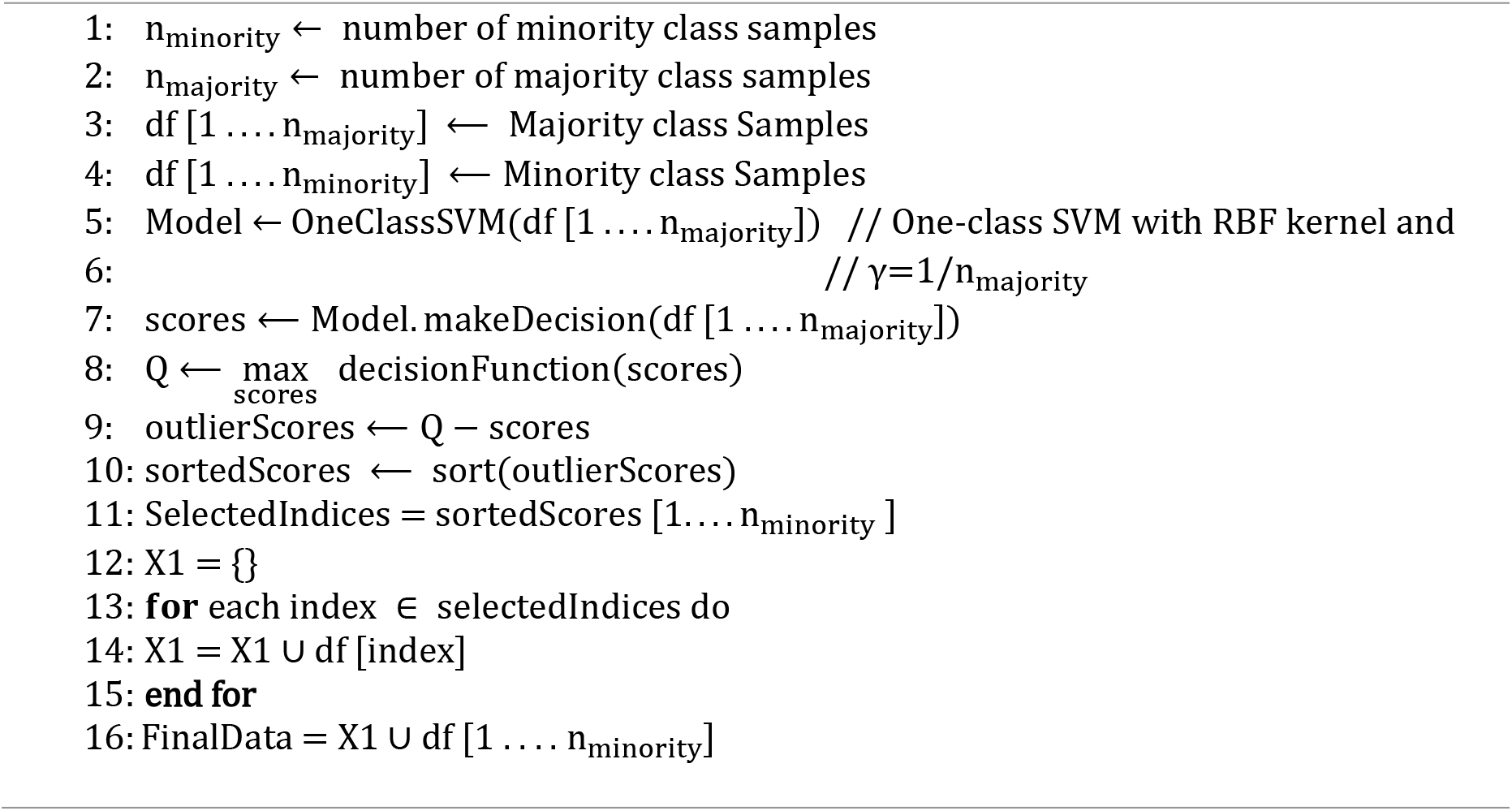

## 5. Feature selection technique

Considering that reducing the number of input features can lead to both reducing the computational cost of modeling and, in some cases, improving the performance of the model. We develop a feature selection algorithm with RF, called FFS-RF. This algorithm was developed and implemented based on the forward feature selection (FFS) technique [66] that coupled with RF to obtain optimal features in DTI. The RF approach [67] is an ensemble method that combines a large number of individual binary decision trees. The performance of the RF model in feature selection was evaluated by a 5-fold CV to construct an effective prediction framework. Forward feature selection is a greedy procedure that iteratively finds the best new feature to add to the set of selected features. The FFS-RF technique starts from the empty feature set and systematically adds features as long as this improves performance, as outlined in Algorithm 2 step by step.

### Algorithm 2: Forward Feature Selection algorithm with RF - (FFS-RF)

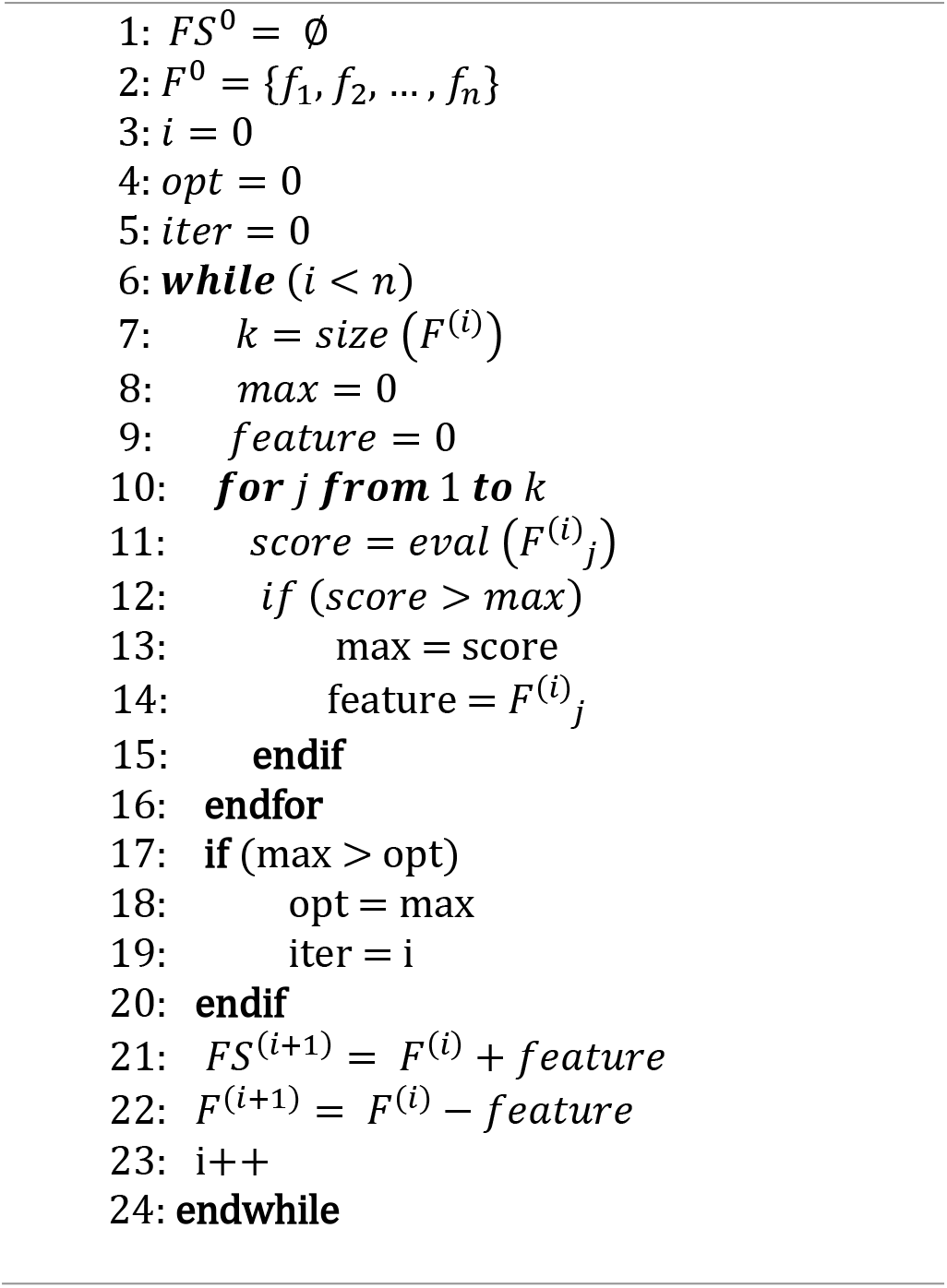

## 6. Results and discussion

In this section, we explain the experimental results of our proposed method in DTI prediction. We implemented all the phases, i.e., features extraction, data balancing, and classifiers of the proposed model in Python language (Python 3.10 version) using the Scikit-learn library. Some of the target descriptors were calculated by the iFeature package [60]and the rest of them were implemented in Python language. OpenBabel Software was used to extract fingerprint descriptors from drugs. All of the implantations were performed on a computer with a processor 2.50 GHz Intel Xeon Gold 5-2670 CPU and 64 GB RAM.

### 6.1 Performance evaluation

Most of the methods in DTI prediction [5, 6, 24, 28] have utilized 5-fold cross validation (CV) to assess the power of the model to generalize. We also use the 5-fold CV to estimate the skill of the model on new data and make a fair comparison with the other state-of-the-art methods. The drug–target datasets were split into 5 subsets where each subset was used as a testing set. In the first iteration, the first subset is used to test the model and the rest are used to train the model. In the second iteration, 2nd subset is used as the testing set while the rest serves as the training set. This process is repeated until each fold of the 5 folds is used as the testing set. Then, the performance is reported as the average of the five validation results for drug-target datasets.

In this study, we perform three types of analyses. First, the importance of feature extraction is discussed. Secondly, we investigate the impact of our balancing technique (One-SVM-US) versus the random undersampling technique on CV results. Finally, the effectiveness of the feature selection method is analyzed.

We used the following evaluation metrics to assess the performance of the proposed model: accuracy (ACC), sensitivity (SEN), specificity (SPE), and F1 Score.

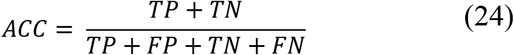

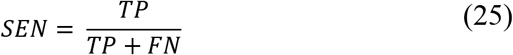

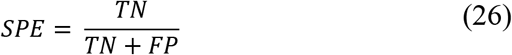

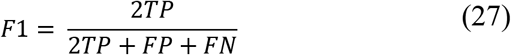

where based on four metrics, namely true positives (TP), false positives (FP), true negatives (TN), and false negatives (FN) are to present an overview of performance. Moreover, we used AUROC (Area Under Receiver Operating Characteristic curve) to show the power of discrimination of the model between the positive class and the negative class. The AUPR (Area Under Precision Recall curve) was also used which would be more informative when there is a high imbalance in the data [68].

### 6.2 The effectiveness of feature groups

We constructed nine different feature groups namely A, B, C, D, E, F, G, H, and I, which all were coupled with drug features to assess the effects of the different sets of features on the performance of the different classifiers including SVM, RF, MLP, and XGBoost. The feature groups have already been reported in Table 2. We also created some subsets from the groups (AB, CD, EF, and GHI), which are given in Table 3. The selection of the best combination can be considered as an optimization problem. Here, we combine feature descriptors based on non-monotonic information and the performance results we get for different classifiers in single feature groups.

We performed experiments to test the effectiveness of the feature groups. In the experiments, we changed the feature groups and applied random undersampling technique to balance datasets. Statistics of the prediction performance for different classifier models are given in Table 4 and Table 5. Focus on the EN dataset, we compared the DTI prediction performance of four different classifiers on nine feature groups and four subsets of them. We also highlighted several possible characteristics that could be considered to select the best classifier in DTI prediction. The results indicated that XGBoost is competitive in predicting interactions. We also made some subsets from single groups namely: AB, CD, EF, and GHI. Two classifiers include MLP and XGBoost had close performance and outperforms other ML methods to predict DTIs.

**Table 4.** Performance of Support Vector Machine, Random Forest, Multilayer perception, and XGBoost classifiers on the gold standard datasets using different feature group combinations and random undersampling technique.

| Dataset | Feature Combination | Classifier | AUROC | AUPR | ACC | SEN | SPE | F1 |
| --- | --- | --- | --- | --- | --- | --- | --- | --- |
| EN | A | SVM | 0.8687 | 0.8642 | 0.7771 | 0.7457 | 0.8085 | 0.7700 |
|  |  | RF | 0.8050 | 0.8042 | 0.7242 | 0.6382 | 0.8103 | 0.6984 |
|  |  | MLP | 0.8939 | 0.8933 | 0.8335 | 0.8635 | 0.8034 | 0.8384 |
|  |  | XGBoost | **0.9253** | **0.9265** | **0.8565** | **0.8447** | **0.8684** | **0.8549** |
|  | B | SVM | 0.9135 | 0.9019 | 0.8429 | 0.8396 | 0.8462 | 0.8425 |
|  |  | RF | 0.8109 | 0.8277 | 0.7378 | 0.6365 | 0.8393 | 0.7085 |
|  |  | MLP | 0.9391 | 0.9378 | 0.8702 | 0.8976 | 0.8427 | 0.8738 |
|  |  | XGBoost | **0.9271** | **0.9254** | **0.8531** | **0.8601** | **0.8462** | **0.8542** |
|  | C | SVM | 0.8457 | 0.8521 | 0.7763 | 0.7457 | 0.8068 | 0.7694 |
|  |  | RF | 0.7832 | 0.7840 | 0.7037 | 0.5939 | 0.8137 | 0.6673 |
|  |  | MLP | 0.8573 | 0.8496 | 0.7925 | 0.8242 | 0.7607 | 0.7990 |
|  |  | XGBoost | **0.9071** | **0.9064** | **0.8386** | **0.8362** | **0.8410** | **0.8383** |
|  | D | SVM | 0.9164 | 0.9042 | 0.8454 | 0.8447 | 0.8462 | 0.8454 |
|  |  | RF | 0.7761 | 0.7949 | 0.7054 | 0.6109 | 0.8000 | 0.6748 |
|  |  | MLP | 0.9385 | 0.9331 | 0.8702 | 0.9027 | 0.8376 | 0.8744 |
|  |  | XGBoost | **0.9307** | **0.9326** | **0.8642** | **0.8584** | **0.8701** | **0.8635** |
|  | E | SVM | 0.8588 | 0.8621 | 0.7788 | 0.7474 | 0.8103 | 0.7718 |
|  |  | RF | 0.8004 | 0.8070 | 0.7208 | 0.6416 | 0.8000 | 0.6969 |
|  |  | MLP | 0.8970 | 0.8954 | 0.8412 | 0.8567 | 0.8256 | 0.8437 |
|  |  | XGBoost | **0.9327** | **0.9348** | **0.8634** | **0.8652** | **0.8615** | **0.8637** |
|  | F | SVM | 0.8777 | 0.8708 | 0.8155 | 0.7628 | 0.8684 | 0.8054 |
|  |  | RF | 0.7972 | 0.8052 | 0.7233 | 0.5751 | 0.8718 | 0.6754 |
|  |  | MLP | 0.9156 | 0.9083 | 0.8352 | 0.9044 | 0.7658 | 0.8460 |
|  |  | XGBoost | **0.9368** | **0.9356** | **0.8745** | **0.8737** | **0.8752** | **0.8745** |
|  | G | SVM | 0.8946 | 0.8972 | 0.8155 | 0.7986 | 0.8325 | 0.8125 |
|  |  | RF | 0.8096 | 0.8176 | 0.7319 | 0.6672 | 0.7966 | 0.7135 |
|  |  | MLP | 0.9254 | 0.9202 | 0.8736 | 0.8754 | 0.8718 | 0.8739 |
|  |  | XGBoost | **0.9317** | **0.9322** | **0.8599** | **0.8652** | **0.8547** | **0.8608** |
|  | H | SVM | 0.8645 | 0.8741 | 0.7797 | 0.7457 | 0.8137 | 0.7721 |
|  |  | RF | 0.8020 | 0.7990 | 0.7293 | 0.6126 | 0.8462 | 0.6937 |
|  |  | MLP | 0.9007 | 0.8999 | 0.8215 | 0.8379 | 0.8051 | 0.8245 |
|  |  | XGBoost | **0.9233** | **0.9216** | **0.8488** | **0.8447** | **0.8530** | **0.8483** |
|  | I | SVM | 0.8931 | 0.8953 | 0.8079 | 0.7833 | 0.8325 | 0.8031 |
|  |  | RF | 0.8132 | 0.8112 | 0.7455 | 0.6485 | 0.8427 | 0.7183 |
|  |  | MLP | 0.9203 | 0.9092 | 0.8676 | 0.8805 | 0.8547 | 0.8694 |
|  |  | XGBoost | **0.9235** | **0.9234** | **0.8497** | **0.8464** | **0.8530** | **0.8493** |

**Table 5.** Performance of Support Vector Machine, Random Forest, Multilayer perception, and XGBoost classifiers on the gold standard datasets using different subsets of feature groups combinations and random undersampling technique.

| Dataset | Feature Combination | Classifier | AUROC | AUPR | ACC | SEN | SPE | F1 |
| --- | --- | --- | --- | --- | --- | --- | --- | --- |
| EN | AB | SVM | 0.9133 | 0.9023 | 0.8429 | 0.8396 | 0.8462 | 0.8425 |
|  |  | RF | 0.8092 | 0.8145 | 0.7404 | 0.6280 | 0.8530 | 0.7077 |
|  |  | MLP | **0.9395** | **0.9373** | **0.8779** | **0.8857** | **0.8701** | **0.8789** |
|  |  | XGBoost | **0.9277** | **0.9290** | **0.8599** | **0.8618** | **0.8581** | **0.8603** |
|  | CD | SVM | 0.9162 | 0.9039 | 0.8454 | 0.8447 | 0.8462 | 0.8454 |
|  |  | RF | 0.7746 | 0.7881 | 0.7096 | 0.6297 | 0.7897 | 0.6846 |
|  |  | MLP | **0.9406** | **0.9387** | **0.8719** | **0.8908** | **0.8530** | **0.8744** |
|  |  | XGBoost | **0.9328** | **0.9379** | **0.8659** | **0.8652** | **0.8667** | **0.8659** |
|  | EF | SVM | 0.8855 | 0.8798 | 0.8155 | 0.7628 | 0.8684 | 0.8054 |
|  |  | RF | 0.7997 | 0.8115 | 0.7190 | 0.5597 | 0.8786 | 0.6660 |
|  |  | MLP | **0.9308** | **0.9315** | **0.8591** | **0.8925** | **0.8256** | **0.8637** |
|  |  | XGBoost | **0.9397** | **0.9411** | **0.8779** | **0.8857** | **0.8701** | **0.8789** |
|  | GHI | SVM | 0.9051 | 0.9020 | 0.8318 | 0.8157 | 0.8479 | 0.8291 |
|  |  | RF | 0.8201 | 0.8351 | 0.7592 | 0.7321 | 0.7863 | 0.7526 |
|  |  | MLP | **0.9278** | **0.9189** | **0.8668** | **0.8805** | **0.8530** | **0.8687** |
|  |  | XGBoost | **0.9288** | **0.9316** | **0.8531** | **0.8618** | **0.8444** | **0.8545** |

### 6.3 The influence of the data balancing techniques

Imbalanced data classification is a significant challenge for predictive modeling. Most of the machine learning algorithms used for classification were designed around the assumption of an equal number of samples for each class. Imbalanced data lead to biased prediction results in ML problems. The drug–target datasets are highly imbalanced. The number of known DTI (positive samples) is significantly smaller than that of unknown DTI (negative samples), which causes to achieve poor performance results of the prediction model. To make balancing in datasets, we used the One-SVM-US technique to build a powerful model. Here, we make experiments to compare the One-SVM-US technique and random undersampling technique to balance datasets in the model. The experimental results are shown in Table 6 and Table 7, which reveals the efficiency of the One-SVM-US algorithm.

**Table 6.** Comparison of prediction results on balanced with Random undersampling and balanced with One-SVM-US in group AB.

| Dataset | Sampling method | AUROC | AUPR | ACC | SEN | SPE | F1 |
| --- | --- | --- | --- | --- | --- | --- | --- |
| EN | Random undersampling | 0.8753 | 0.8625 | 0.8006 | 0.8491 | 0.7887 | 0.8196 |
|  | One-SVM-US | 0.9920 | 0.9975 | 0.9901 | 0.9947 | 0.9967 | 0.9956 |
| GPCR | Random undersampling | 0.7866 | 0.7728 | 0.7354 | 0.7907 | 0.7360 | 0.7727 |
|  | One-SVM-US | 0.9880 | 0.9940 | 0.9732 | 0.9658 | 1.0000 | 0.9826 |
| IC | Random undersampling | 0.8513 | 0.8289 | 0.7873 | 0.8447 | 0.7730 | 0.8233 |
|  | One-SVM-US | 0.9788 | 0.9863 | 0.9543 | 0.9429 | 0.9743 | 0.9565 |
| NR | Random undersampling | 0.6496 | 0.6115 | 0.6556 | 0.9231 | 0.7826 | 0.8000 |
|  | One-SVM-US | 0.9329 | 0.9407 | 0.8611 | 1.0000 | 0.8889 | 0.9474 |

**Table 7.** Comparison of prediction results on balanced with Random undersampling and balanced with One-SVM-US in group EF.

| Dataset | Sampling method | AUROC | AUPR | ACC | SEN | SPE | F1 |
| --- | --- | --- | --- | --- | --- | --- | --- |
| EN | Random undersampling | 0.9024 | 0.9008 | 0.8271 | 0.8737 | 0.8286 | 0.8506 |
|  | One-SVM-US | 0.9910 | 0.9964 | 0.9766 | 0.9647 | 0.9934 | 0.9785 |
| GPCR | Random undersampling | 0.8236 | 0.7762 | 0.7740 | 0.8527 | 0.6800 | 0.7885 |
|  | One-SVM-US | 0.9881 | 0.9928 | 0.9638 | 0.9487 | 0.9562 | 0.9487 |
| IC | Random undersampling | 0.8765 | 0.8527 | 0.8150 | 0.8479 | 0.8191 | 0.8424 |
|  | One-SVM-US | 0.9796 | 0.9880 | 0.9580 | 0.9643 | 0.9807 | 0.9712 |
| NR | Random undersampling | 0.7781 | 0.7029 | 0.7278 | 0.9231 | 0.7391 | 0.7742 |
|  | One-SVM-US | 0.8837 | 0.9033 | 0.8000 | 1.0000 | 0.7778 | 0.9000 |

We observe from Table 6 that the model performance on balanced with Random undersampling and balanced with One-SVM-US in group AB. The results show the significant preference for the AUROC, AUPR, ACC, SEN, SPE, and F1 evaluation metrics by applying One-SVM-US. For the EN dataset, the model achieved AUROC values of 0.9920 in One-SVM-US, and 0.8753 in Random undersampling. In the case of the GPCR dataset, the model obtained AUROC values of 0.9880 and 0.7866, in One-SVM-US and Random undersampling, respectively. For the IC dataset, the model yielded an AUROC of 0.9788 in One-SVM-US and 0.8513 in Random undersampling. Similarly, AUROC values of the model using NR data are 0.9329 in One-SVM-US and 0.6496 in Random undersampling. There is a similar pattern in group EF, which is shown in Table 7. In the case of EN, the prediction results of ACC, SEN, SPE, and F1 on balanced data with One-SVM-US are 0.9901, 0.9947, 0.9967, and 0.9956, which are 0.1895, 0.1456, 0.208, and 0.176 higher than those balanced with Random undersampling, respectively. These prediction results show that the One-SVM-US technique obtains a comparatively advantageous performance. In the case of GPCR, IC, and NR datasets, the ACC, SEN, SPE, and F1 results for balanced data with One-SVM-US and balanced with Random undersampling are in Table 6. The values of these metrics are also shown in Table 7 for group EF. To better analyze the proposed methods, the ROC curves of two data balancing techniques are shown in Fig. 2a–d. These curves demonstrate discriminative ability in group AB, the ROC curve using the One-SVM-US covers the largest area, which is higher than the Random undersampling. The ROC curves of group EF are also shown in Fig. 3a-d, which also cover the larger area in the One-SVM-US technique in comparison with the Random undersampling technique.

**Fig. 2.**
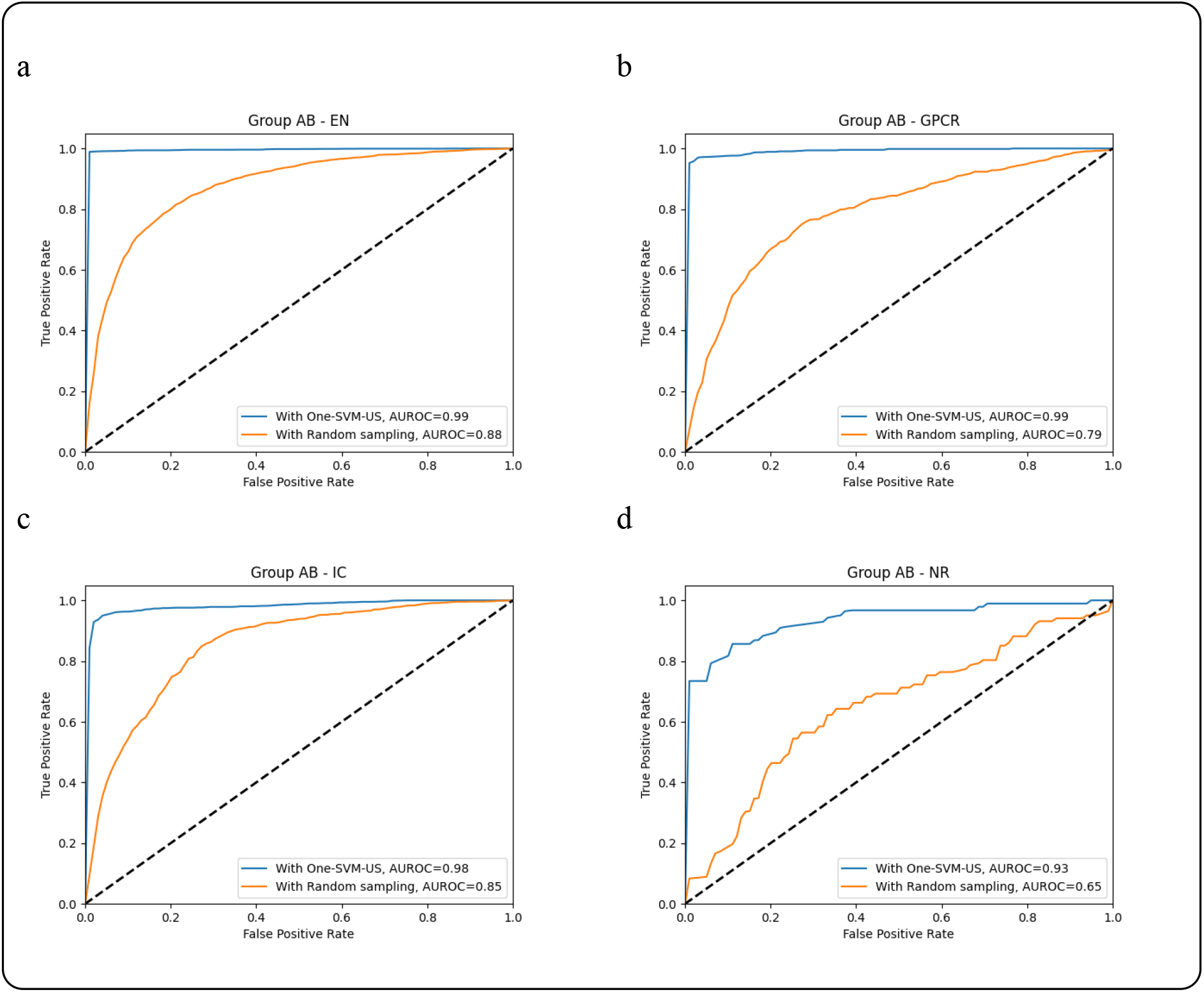
ROC curves of the feature group AB using Random undersampling and One-SVM-US techniques on the datasets: (a) EN, (b) GPCR, (c) IC, and (d) NR.

**Fig. 3.**
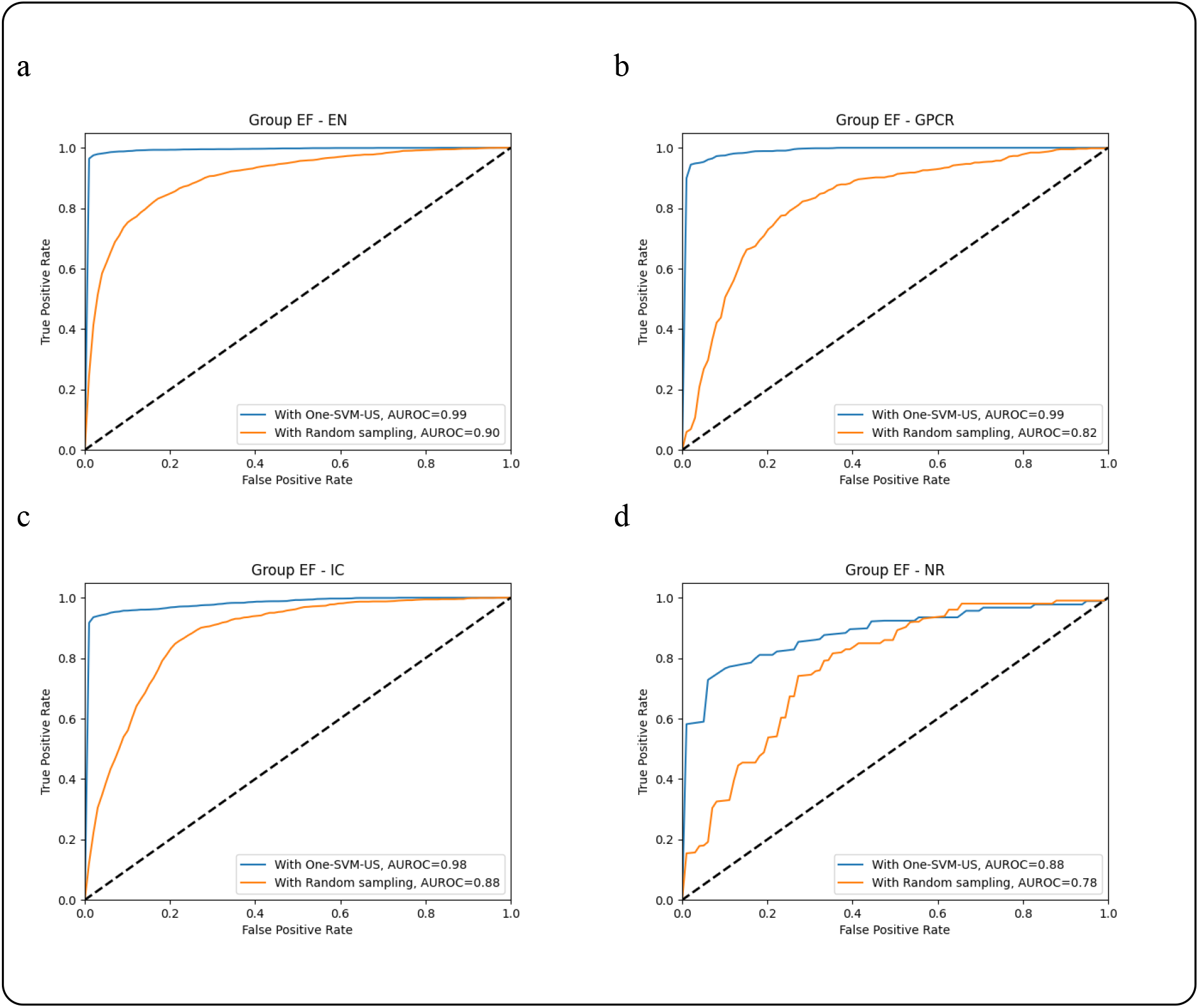
ROC curves of the feature group EF using Random undersampling and One-SVM-US techniques on the datasets: (a) EN, (b) GPCR, (c) IC, and (d) NR.

These results demonstrate that the balanced dataset using One-SVM-US significantly outperforms the balanced dataset using Random undersampling in the case of ROC curves. The accuracy of the XGBoost classifier has been improved after utilizing the One-SVM-US. For all four datasets on the SEN, SPE, and F1 metrics, the results are significantly better in One-SVM-US. Ultimately, One-SVM-US is the efficient method to make balanced datasets to reduce bias and boost the model’s performance.

### 6.4 The effectiveness of feature selection technique

Feature selection is extremely important in ML because it primarily serves as a fundamental technique to direct the use of informative features for a given ML algorithm. Feature selection techniques are especially indispensable in scenarios with many features, which is known as the curse of dimensionality. The solution is to decrease the dimensionality of the feature space via a feature selection method. On the other hand, the more features, the more training time. A feature selection technique by selecting an optimal subset of features reduces computational cost. Various feature selection techniques have been utilized in DTI prediction [1, 6, 63]. The wrapper based methods refer to a category of supervised feature selection methods which uses a model to score different subsets of features to finally select the best one. Forward selection is one of the Wrapper based methods, which starts from a null model with zero features and adds them greedily one at a time to maximize the model performance. Here, we use the FFS-RF algorithm to find the optimal subset and maximize performance. Table 8 indicates the performance results of FFS-RF on the EN dataset in groups AB, CD, EF, and GHI.

**Table 8.**
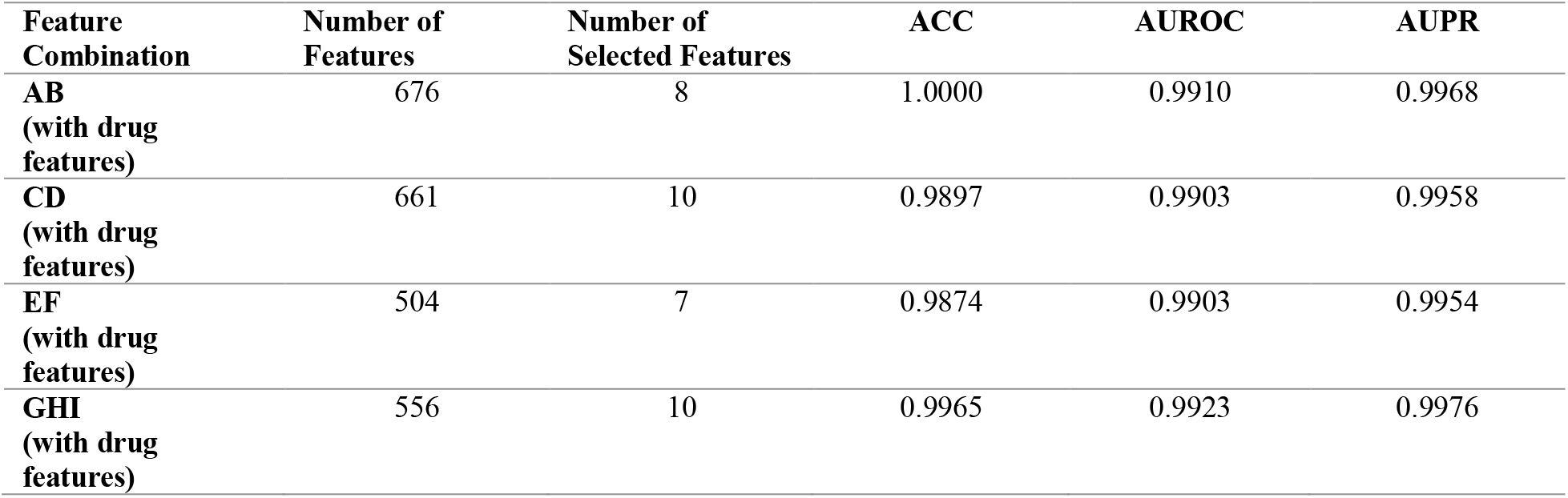
The performance results of FFS-RF on the EN dataset.

Table 8 shows ACC, AUROC, and AUPR metrics of the FFS-RF method which reduces the input features to the model. The worth of the FFS-RF is clearly observable; For the EN dataset, we just use 8 features instead of 676 features in group AB, 10 features instead of 661 features in group CD, 7 features instead of 504 features in group EF, and 10 features instead of 556 features in group GHI. Moreover, the ACC of the FFS-RF method is 100%, 98%, 98%, and 99% in groups AB, CD, EF, and GHI, respectively. The AUROC and AUPR scores are approximately 0.99 in all four groups. In the case of the EN dataset, the feature groups AB and EF had the best and the worst model performance. So, we performed the FFS-RF method on the remaining datasets, i.e. GPCR, IC, and NR for these feature groups. The ACC, AUROC, and AUPR metrics are shown in Table 9 and Table 10 for groups AB and EF, respectively. In group AB, the best feature dimensions selected by FFS-RF are 8, 10, 7, and 10, respectively, which ACC scores are 100%, 99%, 96%, and 97%. The AUROC values of group AB for FFS-RF are 0.9910, 0.9854, 0.9715, and 0.9217. In this group, 0.9968, 0.9924, 0.9769, and 0.9282 are obtained for the AUPR metric. We can see a similar pattern for group EF. Thus, FFS-RF is an effective method to avoid overfitting, improve prediction performance and reduce experimental cost.

**Table 9.** The performance results of FFS-RF on the datasets in group AB.

| Feature Combination | Dataset | Number of Selected Features | ACC | AUROC | AUPR |
| --- | --- | --- | --- | --- | --- |
| AB<br>(with drug features) | EN | 8 | 1.0000 | 0.9910 | 0.9968 |
|  | GPCR | 4 | 0.9921 | 0.9854 | 0.9924 |
|  | IC | 9 | 0.9695 | 0.9715 | 0.9769 |
|  | NR | 4 | 0.9722 | 0.9217 | 0.9282 |

**Table 10.** The performance results of FFS-RF on the datasets in group EF.

| Feature Combination | Dataset | Number of Selected Features | ACC | AUROC | AUPR |
| --- | --- | --- | --- | --- | --- |
| EF<br>(with drug features) | EN | 7 | 0.9874 | 0.9903 | 0.9954 |
|  | GPCR | 10 | 0.9737 | 0.9844 | 0.9893 |
|  | IC | 9 | 0.9762 | 0.9647 | 0.9762 |
|  | NR | 8 | 0.9444 | 0.8993 | 0.9296 |

### 6.5 Selection of predictor model

In this study, we focus on four classifiers: SVM, Random Forest (RF), MLP, and XGBoost. To evaluate these classifier models, we apply Cross Validation (CV) technique to select an appropriate predictor model for our problem. The results of the different predictive models are shown for the EN dataset in group AB in Table 11. To make an obvious comparison of prediction effects, the results are also demonstrated as bar graph for the EN dataset in Fig. 4. Comparison among the prediction results of the EN dataset from Table 11 reveals that the highest results of AUROC, AUPR, ACC, SEN, SPE, and F1 obtained by the XGBoost algorithm are 0.9920, 0.9975, 0.9901, 0.9947, 0.9967, and 0.9956, respectively. The overall prediction ACC of SVM, RF, MLP, and XGBoost is 0.8698, 0.9863, 0.8956, and 0.9901, respectively. The XGBoost ACC is 12%, 0.38%, and 9.45% higher than that obtained by SVM, RF, and MLP classifiers.

**Fig. 4.**
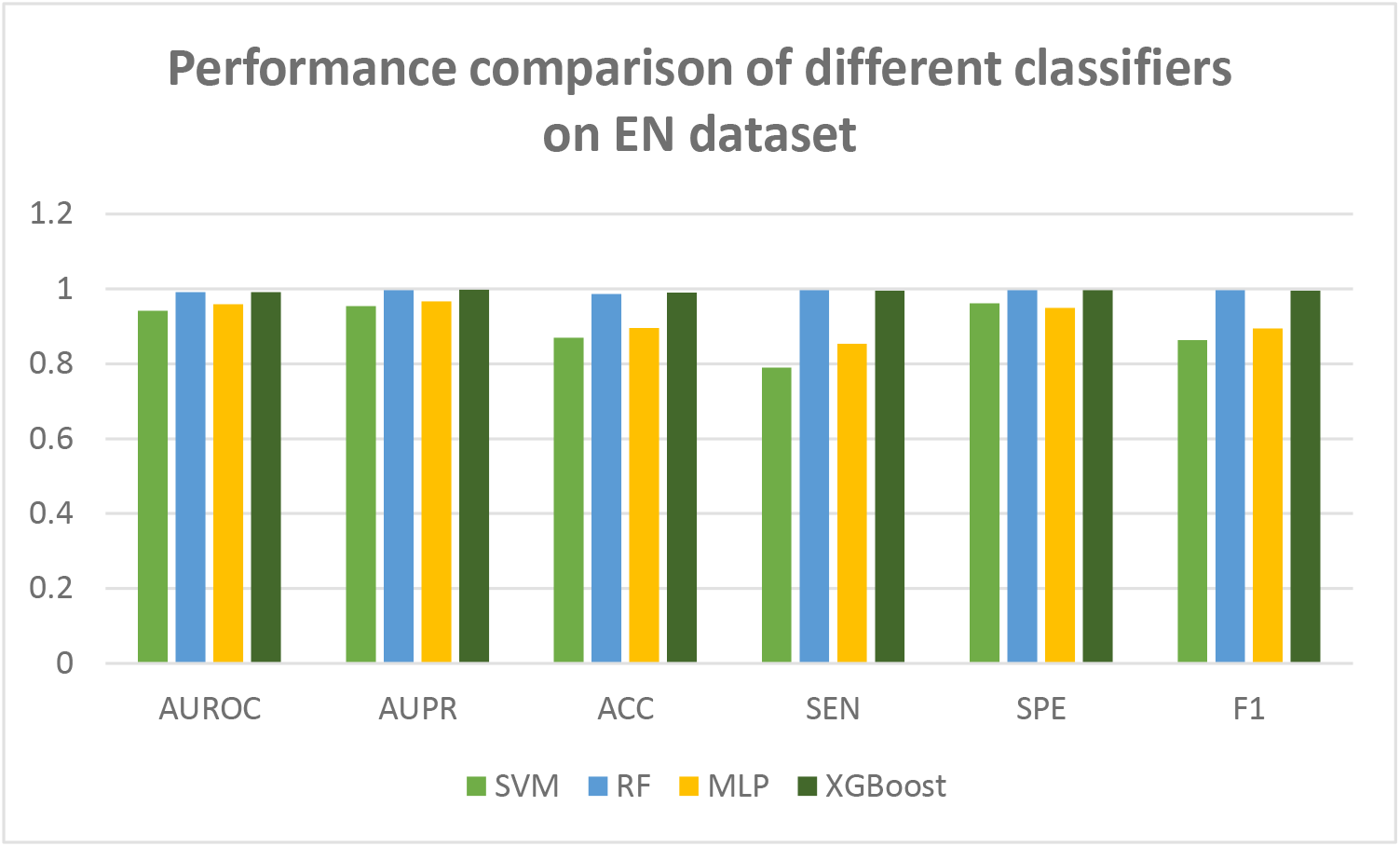
Performance comparison of different feature selection techniques on EN dataset in group AB.

The prediction performance of the XGBoost classifier is premier than the other three classifiers. Therefore, we select the XGBoost classifier as a classification algorithm to predict DTIs.

**Table 11.** The comparison of different feature selection algorithms on EN dataset in group AB

| Classifier | AUROC | AUPR | ACC | SEN | SPE | F1 |
| --- | --- | --- | --- | --- | --- | --- |
| SVM | 0.9417 | 0.9544 | 0.8698 | 0.7898 | 0.962 | 0.8629 |
| RF | 0.991 | 0.9968 | 0.9863 | 0.9965 | 0.9967 | 0.9965 |
| MLP | 0.9586 | 0.9661 | 0.8956 | 0.8534 | 0.9488 | 0.8944 |
| XGBoost | 0.992 | 0.9975 | 0.9901 | 0.9947 | 0.9967 | 0.9956 |

### 6.6 Comparison with other methods

During the last decade, different machine learning frameworks have been proposed to predict DTIs. Some of the proposed methods use feature selection techniques and some of those do not use feature selection. Most of the studies (as well as our approach) have used the dataset proposed by Yamanishi et al. [36] to assess the prediction ability of the proposed methods. To evaluate the effectiveness of our method, we consider six drug–target methods under the AUROC values for the same dataset under the 5-fold CV. In the following, we compare the AUROC of the SRX-DTI model with the other state-of-the-art methods proposed by Mousavian et al. [24], Li et al. [69], Meng et al. [70], Wang et al. [27], Mahmud et al. [63], Wang et al. [26], and Mahmud et al. [6]. The AUROCs generated by these models are listed in Table 12. As seen in the table, the AUROC of the proposed model is superior in comparison with the AUROC of other methods in all the datasets.

**Table 12.**
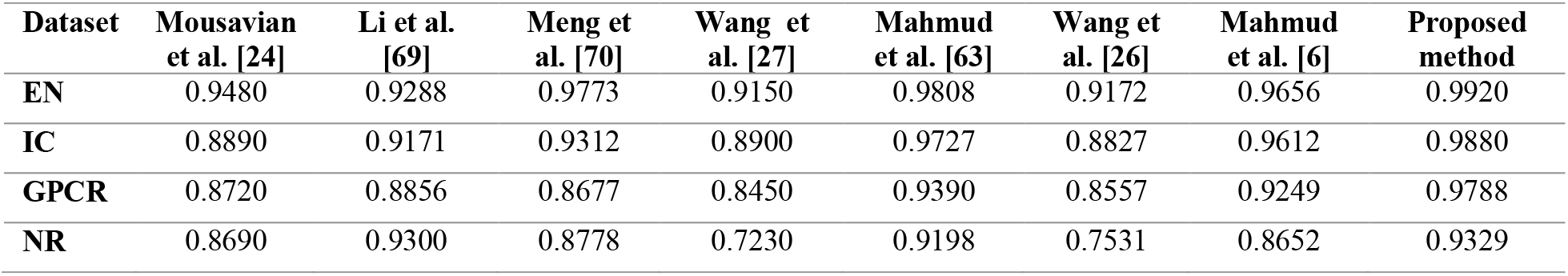
Comparison of proposed model with existing methods on four datasets.

Average AUROC values of SRX-DTI on EN, GPCR, IC, and NR are 0.9920, 0.9880, 0.9788, and 0.9329, respectively. It should be considered that most of the existing models are without a feature selection phase [24, 26, 27, 69, 70]. Training the model with more features can lead to overfitting and reduce the power of generalization in the model. Whereas we can achieve the AUROC of 0.9920 in group AB by using just eight features instead of using all 676 features. This is significantly valuable in terms of computational cost. Moreover, our balancing method superlatively addresses the imbalance problem in the datasets, and feature selection techniques select an optimal subset of features for four datasets. Ultimately, the XGBoost classifier is so scalable that can perform better in comparison with other classifiers for identifying the new DTIs.

## 7. Conclusions

Experimental identification of drug-target interactions is very costly and time-consuming. So, developing computational methods to identify interactions between drugs and target proteins has become a significant step in the drug discovery process for reducing search space to be examined in laboratories. In this work, we proposed a novel framework to predict interactions between drug and target proteins. The novelty of our approach comes in the use of a variety of descriptors for drug and target, implementation of the One-SVM-US technique to address unbalanced data, and development of the FFS-RF algorithm to find an optimal subset of features to reduce computational cost and boost prediction performance. Finally, we compare the performance of four classifiers on the balanced datasets with optimal features. Based on comparison results, the XGBoost classifier is chosen to predict DTIs in our model. The XGBoost classifier is employed to predict DTIs on four benchmark datasets. Therefore, we showed that our robust framework is capable to capture more potent and informative features among massive features. The SRX-DTI model achieved good prediction results, which showed that the proposed method outperforms other methods to predict DTIs.

## Author Contributions

**Hakimeh Khojasteh**: Conceptualization, Methodology, Software, Validation, Formal analysis, Investigation, Writing – original draft, Visualization. **Jamshid Pirgazi**: Conceptualization, Methodology, Formal analysis, Writing – review & editing, Data curation, Supervision, Project administration.

## Acknowledgments

None.

